# Robust and easy-to-use one pot workflow for label free single cell proteomics

**DOI:** 10.1101/2022.10.03.510693

**Authors:** Manuel Matzinger, Elisabeth Müller, Gerhard Dürnberger, Peter Pichler, Karl Mechtler

## Abstract

The analysis of ultra-low input samples or even individual cells is essential to answering a multitude of biomedical questions, but current proteomic workflows are limited in their sensitivity and reproducibility. Here we report a comprehensive workflow that includes optimized strategies for all steps from cell lysis to data analysis. Thanks to convenient to handle 1 μL sample volume and standardized 384 well plates the workflow is easy for even novice users to implement. At the same time, it can be performed semi-automatized using the CellenONE®, which allows for highest reproducibility. To achieve high throughput, ultrashort gradient lengths down to 5 min were tested using advanced μ-pillar columns. Data-dependent acquisition (DDA), wide-window acquisition (WWA) and data-independent acquisition (DIA), and commonly used advanced data-analysis algorithms were benchmarked. Using DDA, 1790 proteins covering a dynamic range of four orders of magnitude were identified in a single cell. Using DIA, proteome coverage increased to more than 2200 proteins identified from single cell level input in a 20-min active gradient. The workflow enabled differentiation of two cell lines, demonstrating its suitability to cellular heterogeneity determination.

## INTRODUCTION

The investigation of individual cells using advanced technologies for cell isolation and analysis of their nucleotide content has revolutionized the current depth of knowledge in the areas of cellular development and behavior.^1–4^ However, cellular identity is driven by its proteome, which is why proteome profiling has become increasingly popular in molecular cell biology research. Reaching single-cell resolution is important because subtle changes in protein expression and turnover can lead to changes in cellular behavior and function. Even clonally identical cells differ from each other depending on their environment and age. Understanding cellular heterogeneity is therefore important to understanding regulatory mechanisms and the development of cells such as stem cells.

To obtain a comprehensive and quantitative proteome profile by classical mass spectrometric techniques requires the bulk analysis of tens of thousands to millions of cells.^5^ Biochemical approaches such as flow cytometry, immunoprecipitations, or immunofluorescence-based techniques help to resolve intercellular networks, but are limited in specificity to approximately a dozen defined proteins.

Single-cell proteomics aims to fill this gap by resolving cellular heterogeneity on global and unbiased levels.^6^ With attomolar detection limits, there is no doubt that mass spectrometry (MS) is sensitive enough to perform this task. The median number of protein copies in an individual mammalian cell is estimated to be 18,000,^7^ meaning that MS identification of at least the most abundant fraction of proteins in a cell is possible. However, reaching sufficient proteome depth remains challenging due to workflow limitations including:

- Sample losses: Due to contact with surfaces and purification processes, the manipulation of cellular content during sample preparation and its transfer into a mass spectrometer can result in losses, leading to significantly lower workflow sensitivity and reproducibility. Techniques recently developed such as SCoPE2^8^, nanoPOTS^9,10^ or the proteoCHIP^11^ minimize sample losses by minimizing handling volumes and automating sample handling, but require specialized equipment.
- Sensitivity: Sample separation and detection of ions in the mass spectrometer must be finetuned and often lack sufficient sensitivity.
- Throughput: To enable definitive statistics, hundreds to thousands of cells must be analyzed. To increase sensitivity and throughput, multiplexed analysis and carrier proteomes are commonly used in single-cell proteomics worflows.^8,10–13^ However, these approaches reduce quantitative accuracy and dynamic range compared to label-free approaches, and the choice of carrier proteome potentially biases the type of peptides identified.^14–16^

Here we present a single-cell analysis workflow that provides a complete set of optimized steps from sample preparation through data analysis. The workflow addresses the limitations associated with current approaches by using standard 384-well plates, which also makes it adoptable by a broader community of scientists. Further, the workflow enables the use of shorter gradients without loss of separating power, increasing throughput, and allowing more robust label-free quantitation (LFQ) strategies. Finally, advanced data acquisition strategies are combined with AI-based data analysis to increase identifications (IDs) and dynamic range.

## RESULTS

### Optimizing cell isolation and digestion to enhance recovery and reproducibility

Aiming for full workflow automatization, ideal cell isolation parameters must be chosen. When using the cellenONE® robot (Cellenion), diameter, and elongation, are the relevant parameters. For this study, these parameters were initially optimized for the selection of individual HeLa cells. To estimate the best set of initial parameters, the distribution of all detected particles was mapped. In line with our expectations^17^, the majority of cells had diameters ranging from 15 to 30 μm. As a first reference, we chose the standard parameters used in our lab before, for the isolation of HeLa cells. We isolated 384 single cells into a 384 well plate. Visual analysis of the images of each isolated cell reveled four different cases: only one cell was isolated per well, more than one cell was isolated per well, the isolated cell appeared apoptotic, or no cell was isolated at all (Figure 1A). Supplemental Figure 1 shows representative examples of each case. Overall, only 88.8% of all wells confidently contained exactly one intact cell. Reducing the diameter range and elongation to a lower value ensured that multiple attached cells are not falsely recognized as single cells and that only intact living cells are isolated by a stricter elongation parameter. Using parameter optimization, the fraction of correctly isolated individual intact cells was eventually increased to 99.7%, which allowed for automation of cell isolation without manually and visually validating images of each isolated particle. The cells were also stained with Hoechst 33342 solution (Thermo Scientific Pierce) to confirm exclusive selection of viable cells.

**Figure 1:**
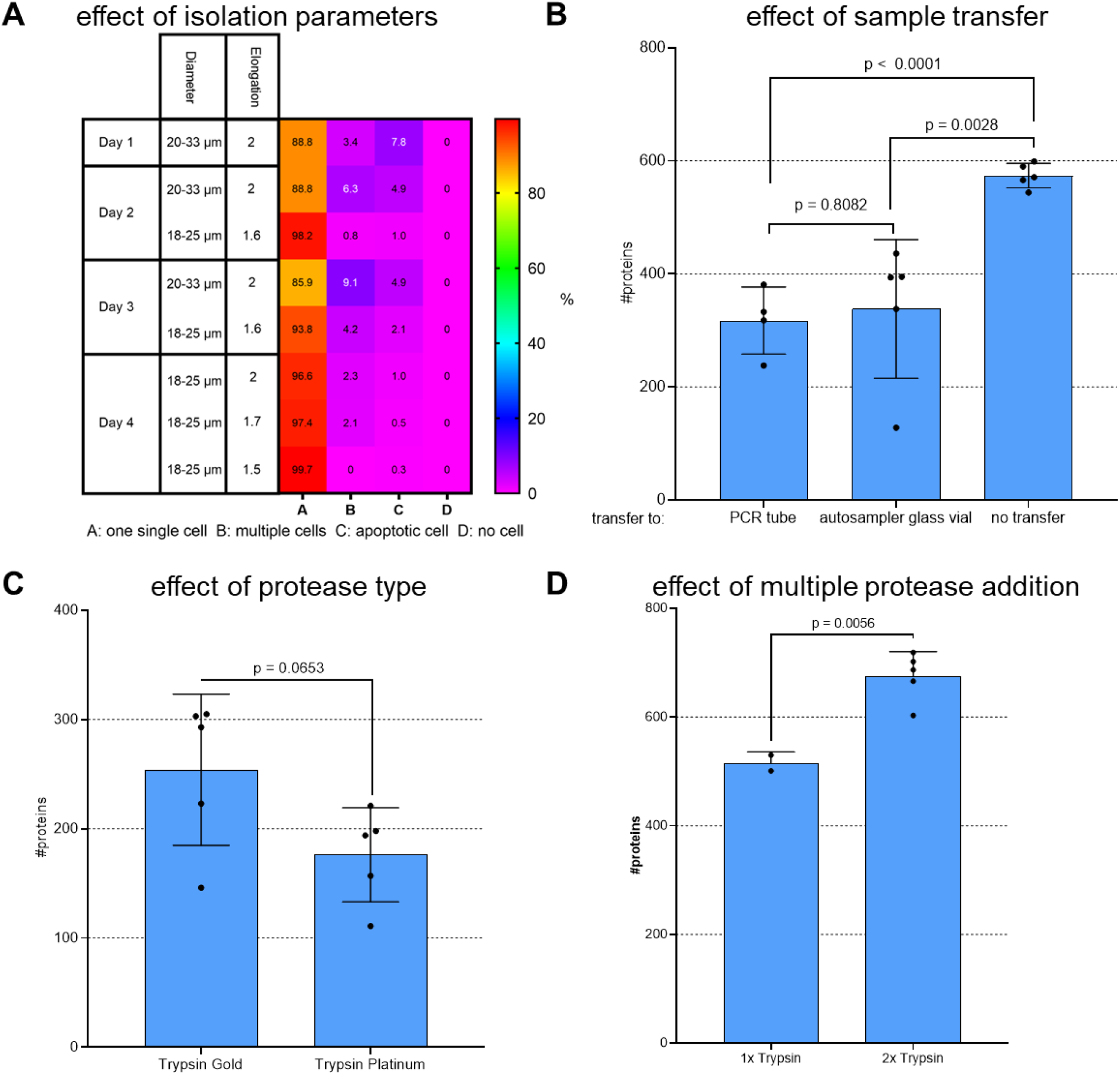
Improving recovery and reproducability by tuning workflow parameters. Individual HeLa cells were isolated into predispensed mastermix using the CellenONE® (**A**): The parameters diameter and elongation were varied on the CellenONE®, numbers indicate the relative amount (%) of cells isolated in the category as indicated, n=384 cells on 4 individual days each. (**B**): Individual HeLa cells were lysed and digested in a 384 well plate and the sample was either injected from there to the LC-MS system or transferred as indicated. (**C**): Cells were digested with Trypsin Gold or Platinum as indicated. (**D**): trypsin was added either once within the mastermix or a second time by addition of 500 nL additional fresh 3 ng/μL trypsin after 30 min. (**B-D**): DDA data was analyzed using CHIMERYS™ at 1% FDR. The bars and error bars show the protein IDs obtained with their standard deviations. An unpaired two tailed Students t-test was performed and resulting p values are depicted above each compared data set, n >4 replicates.

For label free single cell sample preparation, a protocol based on the work of Ctortecka et. al.^11^ was adopted. One μL of master mix containing the detergent DDM for lysis, the enzyme trypsin for digestion and TEAB buffer was predispensed to each well of a 384 well plate followed by isolation of individual HeLa cells into this master mix (for details, see METHODS section). After that, lysis and digestion start with incubation inside the cellenONE® for 2 hours at 50°C and 85% relative humidity. Since sample transfers are prone to loss of hydrophobic peptides, transfer to different tube materials for storage was compared to sample storage and injection directly from the same 384 well plate also used for sample preparation (Figure 1B). Both, transfer to a PEG coated PCR tube, as described by Ctortecka et. al.^11^, as well as transfer to a glass vial lead to obvious losses compared to omitting any sample transfer. In conclusion, all future single cell samples were processed in and injected from a 384 well plate. Next the effect of the used protease (Figure 1C) as well as of multiple protease addition (Figure 1D) was assessed. In detail, Trypsin Platinum and Gold (both Promega) were compared. According to the manufacturer, Trypsin Platinum is free of any detectable nonspecific proteolytic activity which is why an improved overall digestion efficiency is to be expected. Surprisingly Trypsin Gold delivered slightly better results in our single cell setting in terms of protein identifications as well as in terms of observed missed cleavages. Because tryptic activity is highest in the first 30 min of digestion, trypsin was added again after 30 min, which significantly increased the number of identified proteins. Additionally, we tested the protease enhancer ProteaseMAX™ from Promega. On average, we were able to achieve a 17% increase in the proteins found in the single cell samples by adding the enhancer to our trypsin digestion (Supplemental Figure 4). Therefore, all subsequent experiments were performed by adding ProteaseMAX™ to the mastermix and Trypsin Gold twice. The detailed method steps used to perform the cell isolation, sample transfer, and digestion experiments are provided in METHODS.

### Optimizing sample preparation to boost protein IDs

Though using 1 μL instead of nanoliter-range volumes makes sample handling more convenient, it increases wetted surface area and the chance that peptides will stick to the tube walls. With this in mind, we attempted to keep peptides solubilized by supplementing 5% DMSO. Use of DMSO was tested on diluted bulk digests at a concentration of 250 pg / 3.5 μL, which was the same as the concentration expected in the real single-cell samples under ideal conditions. Demonstrated by the additional protein IDs obtained, DMSO reproducibly and significantly increased recovery (Figure 2A). The additional peptides found in the DMSO-containing samples predominantly eluted towards the end of the gradient (Figure 2B), supporting the conclusion that DMSO supplementation improves the solubility of hydrophobic peptides. In addition, the vast majority of 80% of all peptide-spectrum matches (PSMs) found in a representative replicate analysis performed without using DMSO were ultimately found when DMSO was added (Figure 2C). Those not found could be due to the inherent run-to-run variability of the stochastic DDA method used.

**Figure 2:**
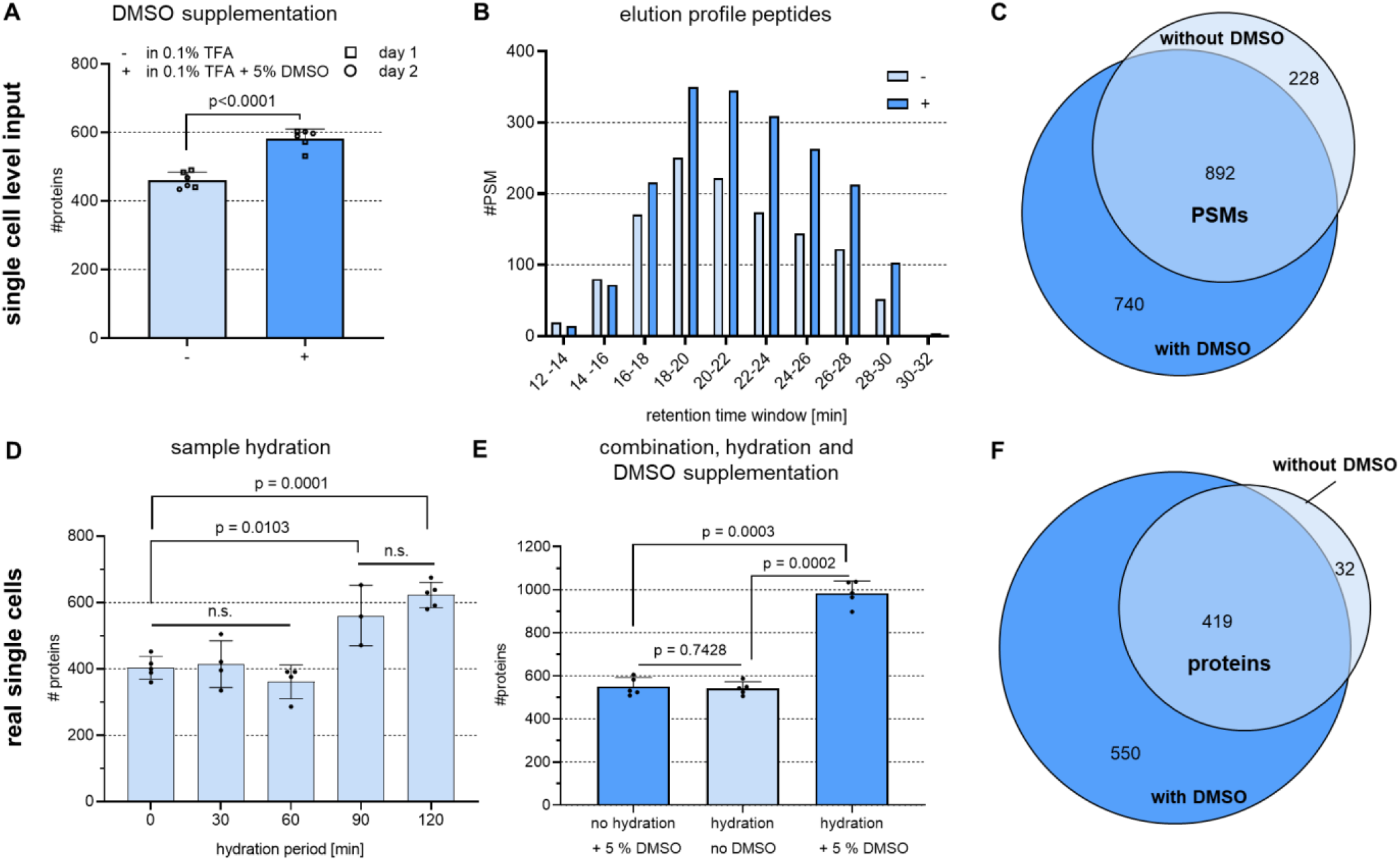
DMSO supplementation and sample hydration improved recovery of hydrophobic peptides and boosted IDs. All data were acquired using DDA and then analyzed using the CHIMERYS™ algorithm with 1% FDR at the PSM and protein levels. (**A**): 250 pg HeLa bulk digest dissolved at a concentration of 71 ng/μL in the presence or absence of 5% DMSO prior to injection into the LC-MS. The bars and error bars show the resulting protein IDs and their standard deviations. To estimate significance, an unpaired two-tailed Students t-test was performed. The resulting p values are shown above each compared data set, n = 6 technical replicates measured on 2 different days. (**B**): Data from a representative replicate of the DMSO tests performed in panel A. The bars present the PSM matches in each retention-time window. (**C**): Venn diagram showing the common PSMs from the protein IDs shown in panel B. (**D**): Individual HeLa cells were lysed and digested at 50 °C for 2 h while the sample was hydrated with automated water supplementation for the time periods shown. The bars and error bars show the resulting protein IDs and their standard deviations. A one-way ANOVA with Tukey’s multiple comparison post-test was performed, n ≥ 3 replicates. (**E**): Individual HeLa cells were kept hydrated for 2 h or dried upon evaporation during digestion in the presence or absence of DMSO in the final storage solution. The bars and error bars show the resulting protein IDs with their standard deviations. An unpaired two-tailed Students t-test was performed and resulting p values are shown above each compared data set, n = 5 replicates (**F**): Venn diagram showing the common protein IDs from a representative replicate comparing hydration and no DMSO with hydration plus 5% DMSO from the data shown in panel E.

To minimize adsorptive sample losses also during single cell sample preparation we further aimed to hinder sample drying by automated addition of water during digestion at 50 °C within the CellenONE®. By that the digestion mix was kept hydrated over a period from 0 - 120 min until the end of incubation. In case of no hydration task performed, the sample was reproducibly dried after ∼30 min, which is why 500 nL water were added every 15 min to keep the volume at the same level. Our results suggest that an elongated hydration allows for longer active digestion and lowered sample adsorptive sample loss as indicated by more proteins identified when hydrating at least over the first 90 min of the digestion process (Figure 2D). Analyzing single cells that were either kept hydrated during digestion or supplemented with DMSO for storage lead to roughly 500 protein IDs. Combining both boosting strategies results in more than 1000 average proteins identified from a single HeLa cell using DDA and analyzing data with CHIMERYS™ on 1 % FDR at peptide and protein level (Figure 2E). This effect is even more distinct on peptide level, where an average of 1883 peptides was found without DMSO of hydration compared to 4625 with hydration and DMSO, which corresponds to a boost by 146 %. Notably, adding DMSO enabled identification of 550 additional proteins, with 93% of all proteins previously identified in the absence of DMSO found (Figure 2F). All data were acquired using DDA and then analyzed using the CHIMERYS™ intelligent search algorithm (MSAID^®^) with 1% FDR at the PSM and protein levels. The detailed method steps used to perform the sample preparation experiments are provided in METHODS.

### Sample preparation workflow key points

To ensure a robust, reproducible, and automated single-cell workflow, single-cell isolation was performed using the cellenONE® picolitre dispensing robot. Using a 384-well plate and easy-to-handle 1 μL volumes allows laboratories to implement the workflow without need for expensive, specialized, ultralow-flow equipment and therefore also in combination with other cell isolation machines as a FACS device commonly available in many lab environments. Constant hydration was automatically performed by the cellenONE® but can be replaced by manual pipetting. Our optimized workflow is sketched in Figure 3 and detailed within the methods section.

**Figure 3:**
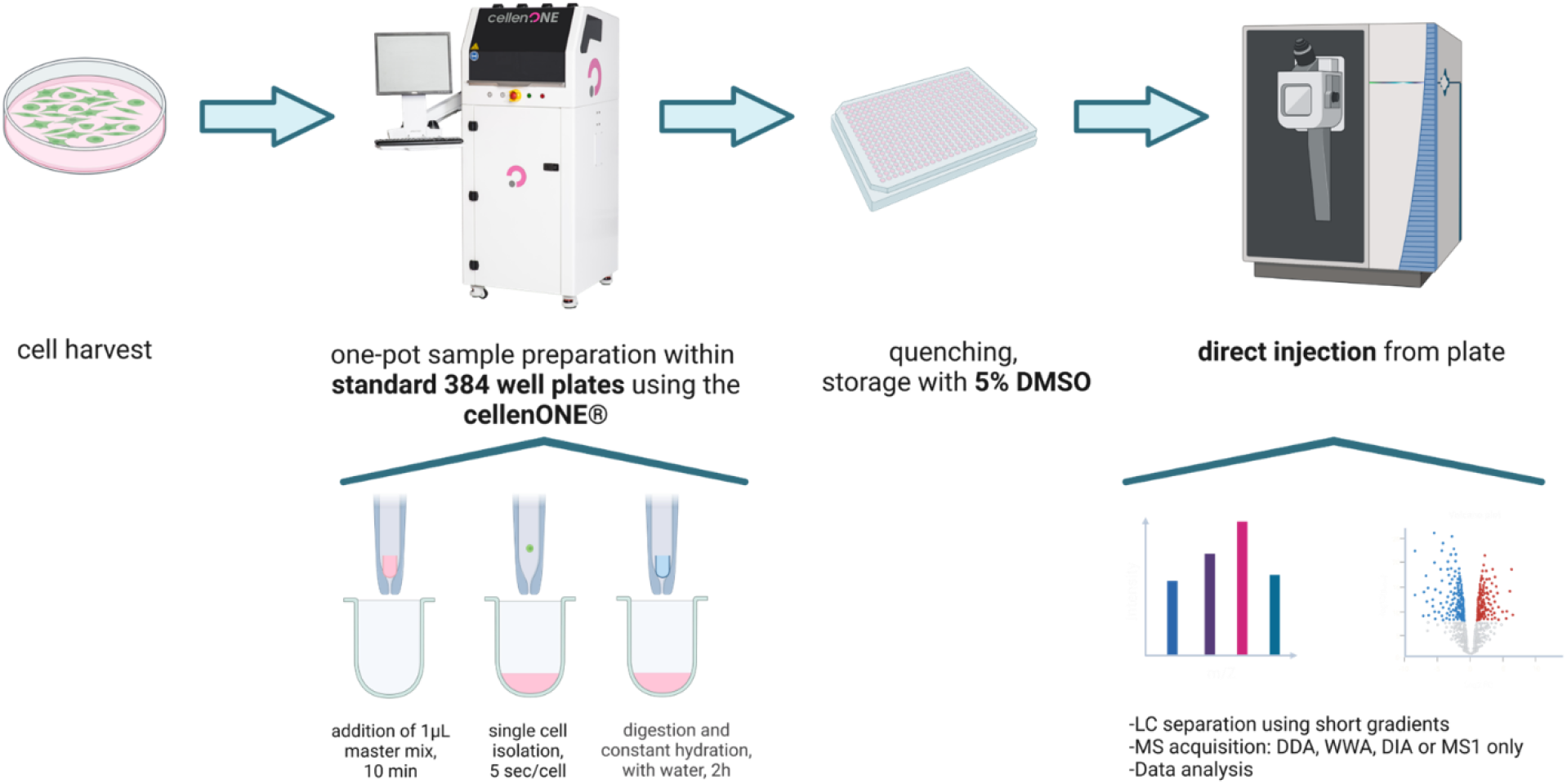
Overview label free, one-pot, single cell sample processing workflow. Cells are harvested followed by lysis and digestion within the CellenONE® in a 384 well plate that is also used for direct placement into the autosampler of the HPLC-MS. Figure created with Biorender.

### Optimizing LC-MS data acquisition and analysis to maximize IDs

In parallel to parameter tuning for a loss free and robust single cell sample preparation, we aimed to improve proteome coverage by optimizing data acquisition and analysis as well. In a first step we compared two column types for their chromatographic performance: A classical packed bed column (CSH C18 Column, 130Å, 1.7 μm, 75 μm X 250 mm, Waters) that was previously used in our lab and a μPAC™ column (brick shape pillars, 5.5 cm, prototype column, Thermo Fisher Scientific) which we recently successfully tested for low input amounts as well as short gradients.^18^ Although the μPAC™ column is only 5.5 cm long, its internal flow path is roughly 50 cm leading to a sufficient surface area enabling powerful separation. The median FWHM peak width obtained using the packed bed column was 3.84 sec vs 4.77 sec for the μPAC™. Although peak withs seem slightly wider, the μPAC™ column clearly and reproducibly outperformed its conventional counterpart in ID numbers (Figure 4C), which is why we used it for following optimizations as well.

**Figure 4:**
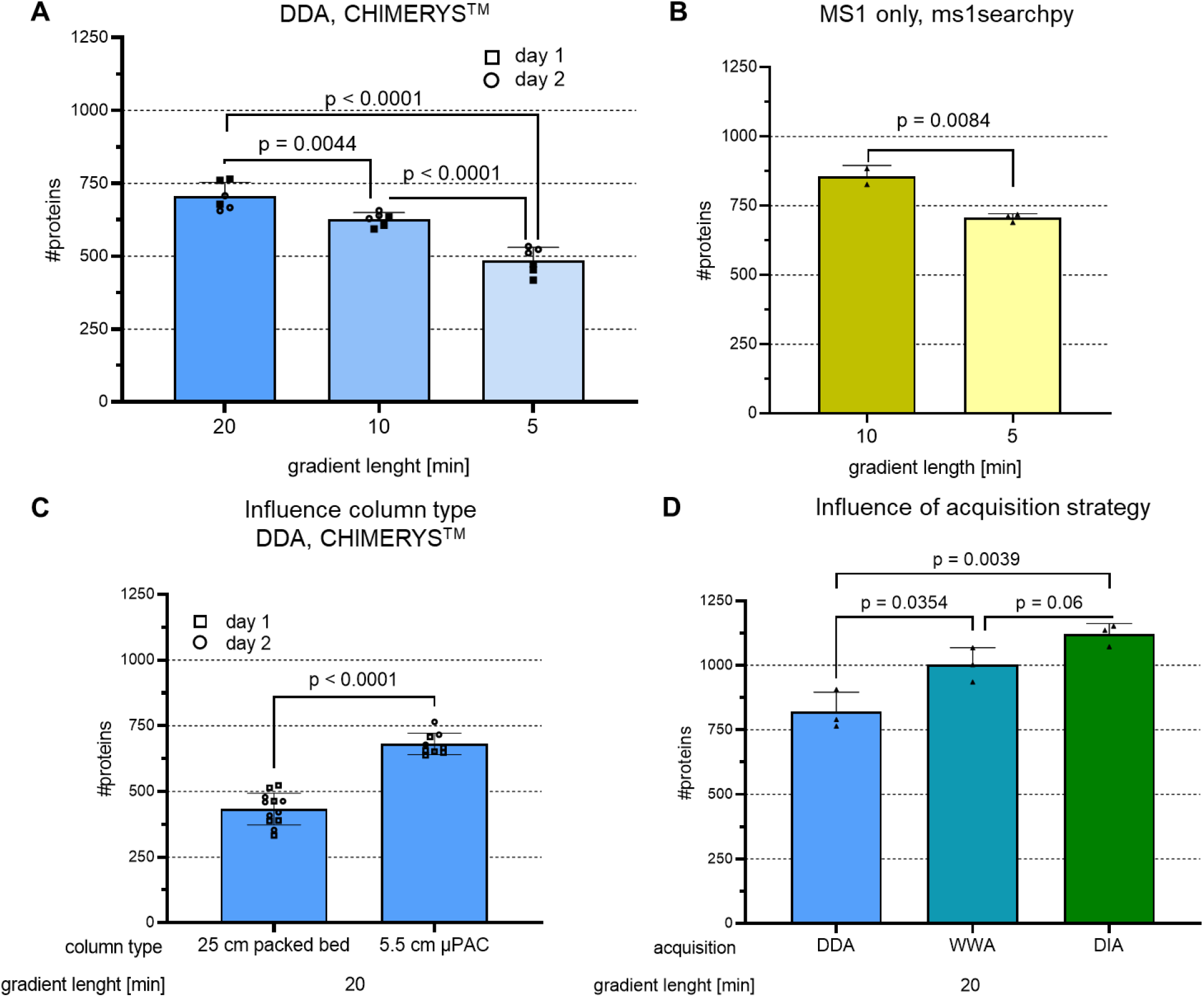
Influence of gradient length, column type and data acquisition strategy on protein IDs of single cell level inputs. 250 pg HeLa bulk digest was used for injection into LC-MS each, bars represent protein IDs on 1 % FDR level, error bars indicate standard deviations. To estimate significance, an unpaired two tailed Students t-test was performed and resulting p values are depicted above each compared data set. (**A**): Peptides were separated on a 5.5 cm μPAC™ column, data was acquired using DDA and analyzed using CHIMERYS™, n = 6 technical replicates measured on 2 different days. (**B**): As A but data analysis was performed using ms1searchpy^19^, n= 3 technical replicates. (**C**): Peptides were separated either on a 25 cm packed bed column or on a 5.5 cm μPAC™ column, data was acquired using DDA and analyzed using CHIMERYS™, n = 12 technical replicates measured on 2 different days. (**D**): As A but using either DDA (2 m/z isolation width), WWA^18^ (12 m/z isolation width) or DIA (variable windows 8 – 25 m/z isolation width9 as acquisition strategy. DDA and WWA data were analyzed using CHIMERYS, DIA data was analyzed using Spectronaut® in directDIA mode, n = 3 technical replicates.

Minimizing runtimes using short gradients offers the potential advantages of increased throughput and sensitivity due to sharper peak shapes. Unfortunately, reducing an already short 20-min active gradient to 5 min lowers protein IDs from 705 to 485 proteins or by about 30% (Figure 4A). The resulting peaks are sharp with a median FWHM of 4.77 and 2.9 sec for the 20- and 5-min gradients, respectively, which is a prerequisite for high sensitivity. However, both the 20- and the 5-min gradients make it difficult for a Thermo Scientific™ Orbitrap™ mass analyzer to record enough MS2 spectra to obtain the desired IDs. To address the need for increased throughput while achieving sufficient proteome coverage, the mass spectrometer duty cycle can be reduced by omitting requirements for fragment spectra. The first successful attempt to perform proteome-wide analyses using very short gradients of only 5 min was carried out by Ivanov and coworkers in 2020^19,20^. Their approach was adopted for the single-cell workflow experiments described here. The 5.5 cm μPAC™ HPLC column was used, which, due to its low backpressure, allowed for fast loading and re-equilibration at 1 μL/min in a total run time of 7.4 min. For short gradients, the MS1-only approach outperformed standard DDA. Roughly the same number of proteins were identified in the 5-min and 20-min active gradients (Figure 4A and B). Of those proteins found in the 20-min DDA run, approximately 70% were identified in the 5-min MS1 run, which is a comparable to published results for higher input samples^19^ (Supplemental Figure 2).

Different mass spectrometer data-acquisition strategies were then tested using the 20-min active gradient (Figure 4D). The proteome depth obtained using DDA was improved by widening the precursor isolation window from 2 to 12 *m/z*. Using such a WWA^18^ on purpose generated chimeric spectra and led to the identification of more than one peptide from a single spectra when the CHIMERYS™ intelligent search algorithm was used. Variable DIA windows of up to 25 *m/z* for co-fragmentation of ions significantly increased protein IDs to more than 1100 from 250 pg HeLa.

Due to the superiority of the DIA we investigated this strategy in more detail. Figure 5A shows the advantage of using variable rather than fixed windows. Using variable windows reduces duty cycle because broader windows are used in *m/z* regions where few ions are expected. The mass spectrometer parameters used are provided in **Error! Reference source not found**.. Similar to the results obtained for the DDA experiments, protein IDs were increased using DIA, in this case by about a factor of two, when the μPAC™ HPLC column was used instead of the packed-bed column (Figure 5A versus Figure 5B). In both experiments, the same method and gradient were used (dark green bar, 20-min gradient, variable windows). To dig deeper into the proteome and improve data completeness across replicates, matching between runs was performed (Figure 5B, no reference library). Across three replicates, data completeness increased to nearly 100% with almost 1500 proteins identified from 250 pg sample using a directDIA search. Enriching the directDIA search data with data obtained from higher input samples of 1 or 10 ng improved matching, boosting protein IDs to more than 2000 and 2200, respectively. Enlarging the library size however led to identifications not possible in all replicates (lower recovery), reducing data completeness to approximately 80% when the largest library was used. Of note, these results are in line with our observations made on a timsTOF with DIA-PASEF and using a library for enrichment using single cells (Pichler et. al., manuscript in preparation).

**Figure 5:**
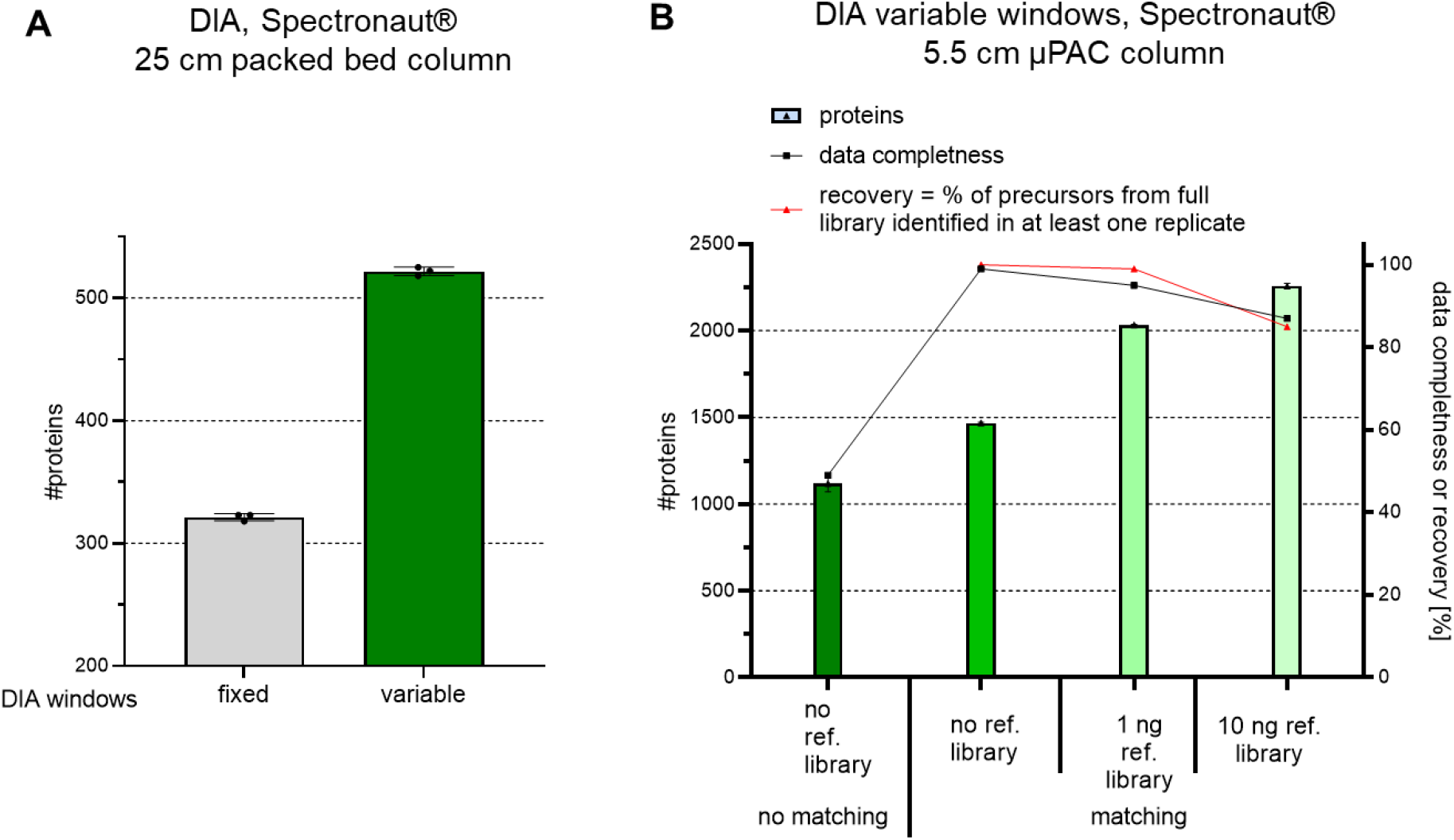
Benchmarking DIA methods and enrichment of searches to boost protein IDs of single cell level inputs. 250 pg HeLa bulk digest was prepared for single-cell LC-MS analysis for each experiment, bars and error bars show the protein IDs at the 1% FDR level with their standard deviations. (**A**): Peptides were separated on a 25 cm packed-bed column. Data were acquired using a fixed window size of 8 m/z or variable windows ranging from 8 – 25 m/z, and then analyzed using Spectronaut®, with n = 3 technical replicates. (**B**): Peptides were separated on a 5.5 cm μPAC™ HPLC column. Data were acquired using variable windows ranging from 8 – 25 m/z, and then analyzed using Spectronaut®, either separately (no matching) or together (matching between runs), with or without enrichment using data from higher input experiments, with n = 3 technical replicates each.

Using match between runs (MBR) also improved the results for DDA-based single-cell analysis. Five replicates of one individual HeLa cell each were analyzed using four of the commonly used data analysis algorithms (Figure 6A). The MS Amanda^21^ and CHIMERYS™ algorithms were each evaluated in combination with apQuant^22^ as nodes within Proteome Discoverer™ software (Thermo Scientific). Using MBR yielded more than 100 additional protein IDs in each case. A similar effect was observed when using MSFragger^23^ database search tool and IonQuant^24^ data analysis. In contrast, the number of protein IDs was increased by only 5% when using MBR with SpectroMine™ (Bionosys). To improve sensitivity, five replicates of 40 HeLa cells were analyzed together with the single cell runs to provide more potential matches across files (Figure 6B). As a result, the matching library was increased to 7327 peptides and 1201 proteins for the MS Amanda searches, and to 16229 peptides and 2955 proteins for the CHIMERYS™ searches. The resulting protein identifications in single cells increased by up to 71%, or to 999 and 1791 average proteins for the Amanda and CHIMERYS™ searches, respectively. In contrast, using SpectroMine™ or MS Fragger in combination with the additional 40 cell files, did not significantly change the resulting protein IDs. With or without MBR, CHIMERYS™ outperformed all the other search engines tested, making it the preferred choice for DDA and WWA searches in this study.

**Figure 6:**
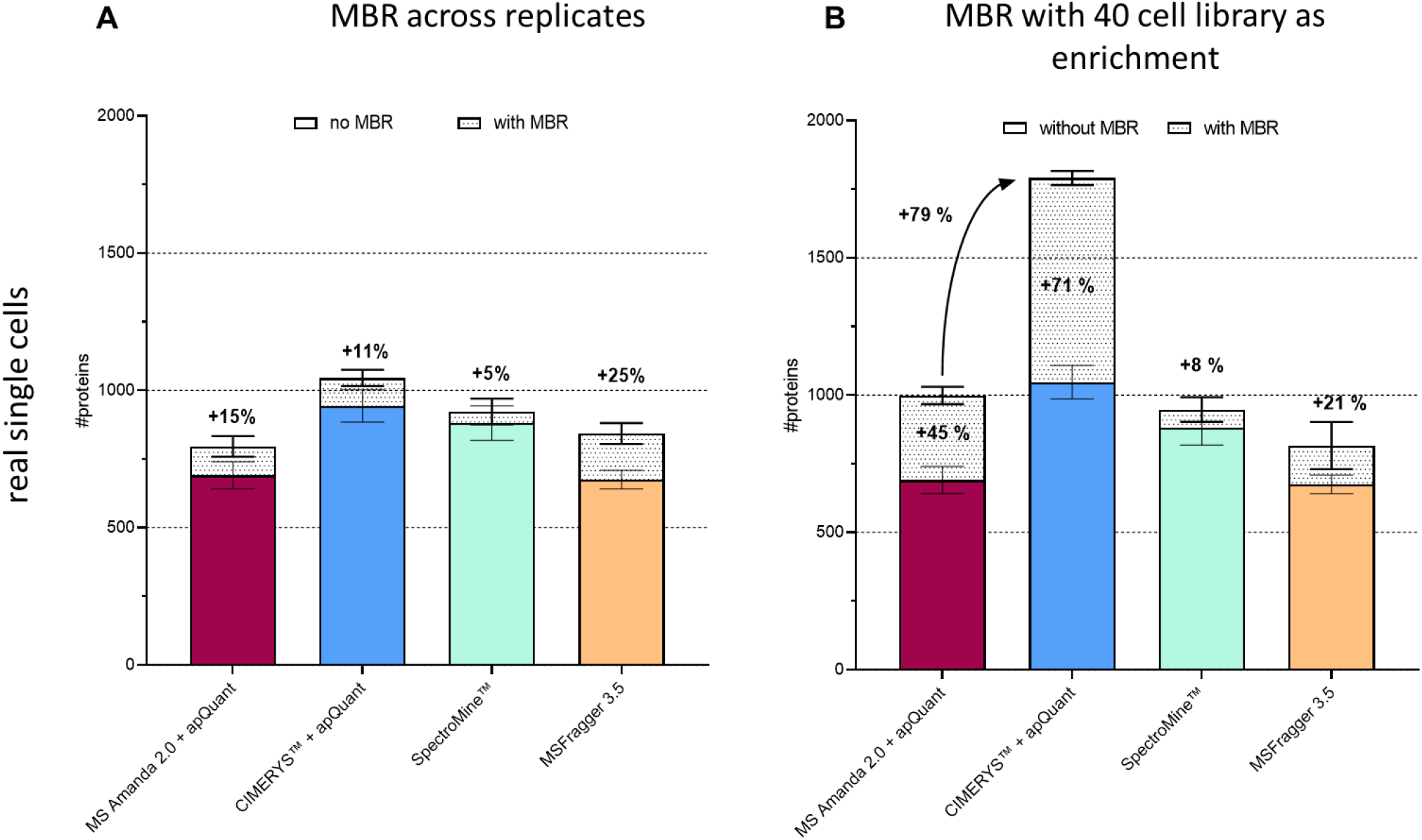
Benchmark of different data analysis tools and effect of enrichment for DDA runs. Individual HeLa cells were processed using the optimized workflow. Peptides were separated using a 20-min active gradient and data were acquired using a DDA method with a 2 m/z isolation width. (**A**): Data analysis was performed using each evaluated software tool at 1% FDR on the protein level, either file-by-file or with MBR, across all n = 5 replicates. (**B**): As A but data processed together with five runs of 40 HeLa cells each to enrich matching.

### Proof of principle studies Proteome depth

Reaching sufficient proteomic depth is essential for single-cell workflows that aim to investigate cellular heterogeneity. To estimate the sensitivity of the workflow presented here, all proteins identified from a single-cell run that yielded more than 1000 proteins were plotted into a histogram showing protein IDs obtained from a high-input sample with more than 8500 proteins quantified. As shown in Figure 7A, the single-cell analysis results predominantly covered proteins that are most abundant in the proteome. Nevertheless, the identified proteins ranged over more than four orders of magnitude in abundance, which is sufficient to allow investigation of differences in protein abundance. This broad dynamic range covered by the single-cell run was estimated by plotting all PSM intensities (Figure 7B). Notably, the most abundant hits were contaminants introduced during sample preparation; mostly keratins and trypsin which is added in excess for digestion.

**Figure 7:**
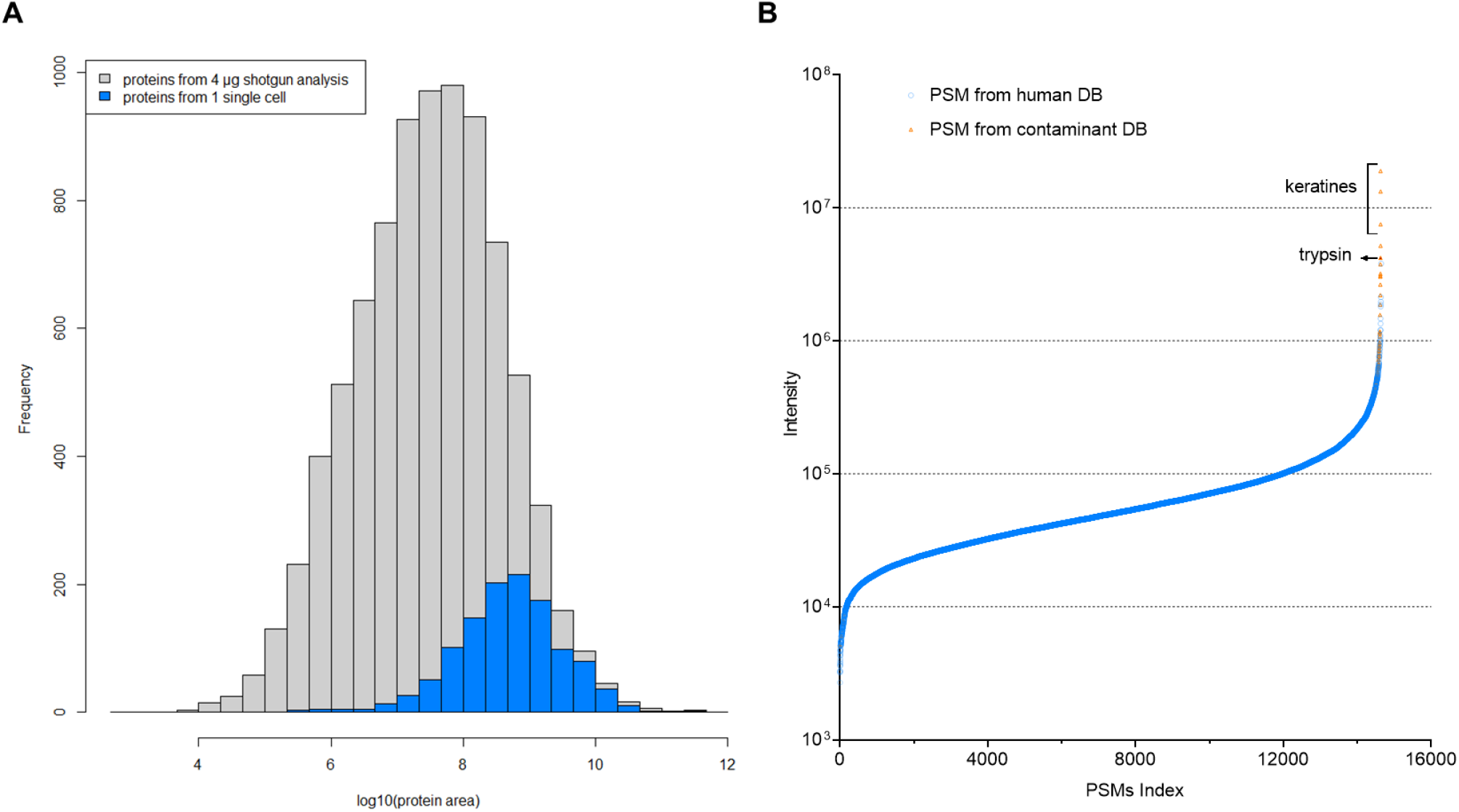
Proteome depth reached in single-cell experiments, using DDA and label-free strategies. Data from a single HeLa cell obtained using the optimized workflow, acquired in DDA mode and analyzed using CHIMERYS™. (**A**): Proteins, excluding contaminants, identified from a single cell, plotted on the abundance distribution of all proteins identified in a 4 h shotgun run from 4 μg HeLa digest. LFQ was performed using apQuant. (**B**): PSMs identified in a single-cell analysis, indexed based on their precursor intensity. PSM matches originating from the contaminant database (DB) are highlighted in orange.

To ensure that those IDs that are of low precursor signal intensity and of a spectrum quality that is just accepted by the 1% FDR threshold, we performed an additional analysis including an entrapment of the yest proteome into the database. We found, that 1.5 % of all peptides and 1.3 % of all proteins in the result file were originating from the yeast proteome, which indicates a proper overall FDR calculation. Furthermore, those PSMs identified from yeast are not accumulated in the low abundant fraction but evenly distributed across the whole intensity range. This indicates that not only noise is reported in the low abundant fraction and the achieved dynamic range is truly in the range of 4 orders of magnitude (Supplemental Figure 3).

### Application of optimal workflow parameters to single-cell analysis of two human cell lines – proof of principle study

To evaluate the optimized workflow, individual HeLa and K652 cells were isolated, processed and analyzed using the optimized sample preparation conditions. Cells were kept hydrated, trypsin was added twice, and DMSO was supplemented for storage. Peptides were separated using a 20-min active gradient on a 5.5 cm μPAC™ HPLC column and data were acquired using a DIA method with variable windows. Even with relatively few replicates, clear separation of cell types was obtained based on their proteome abundance variation (Figure 8A), indicating that the workflow is sensitive enough to enable investigation of cellular heterogeneity. This is mirrored when visualizing this data using a dendrogram (Supplemental Figure 5) where we also included data from 40 cells measured (used for enrichment) or from a no cell control sample. On average, 1208 and 1139 proteins were identified from a single HeLa or K562 cell, respectively (Figure 8B). Matching to measurements of 40 HeLa or K562 cells boosted IDs to 1543 and 1404 proteins respectively.

**Figure 8:**
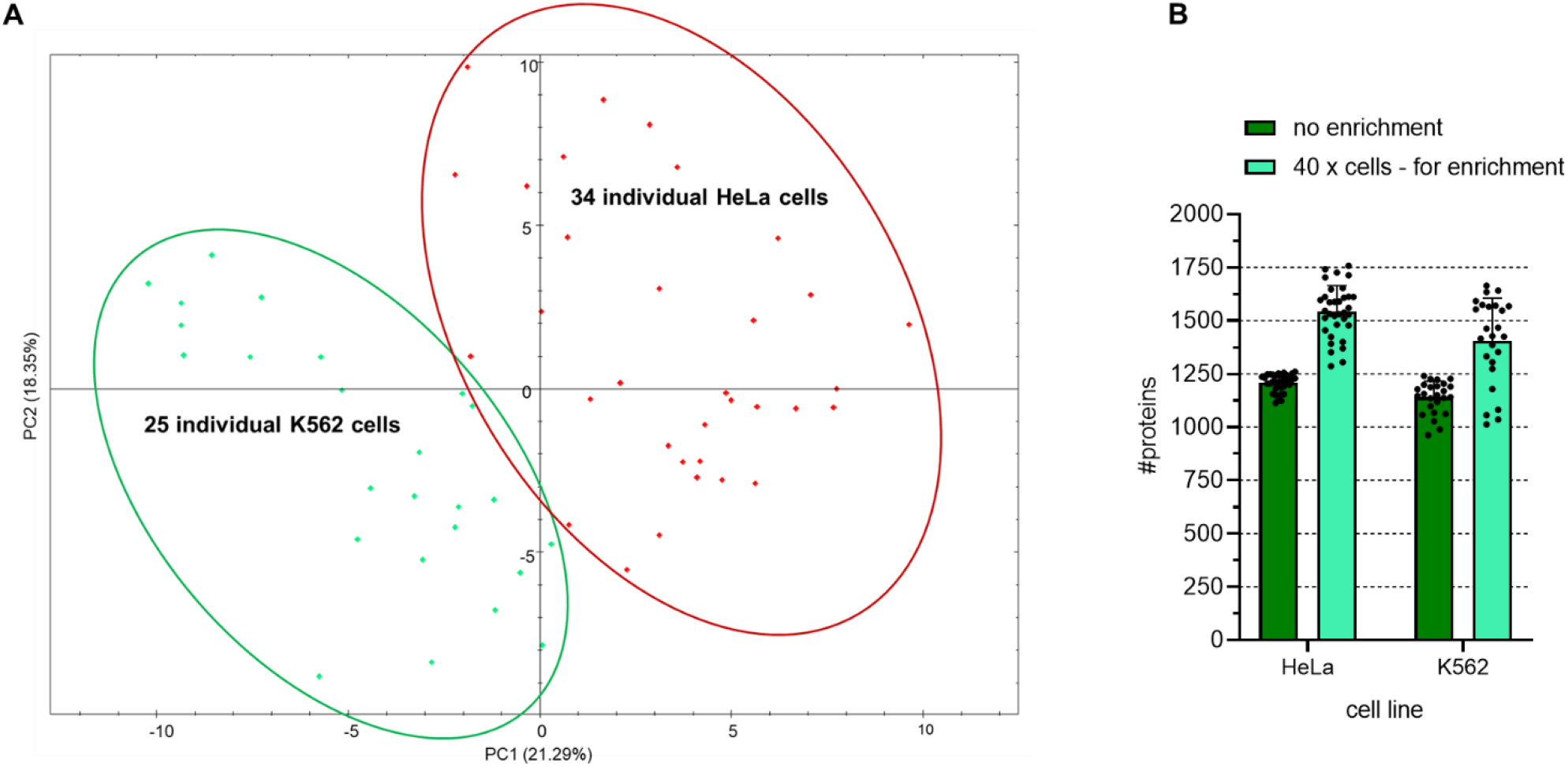
Application of optimal sample preparation, data acquisition, and data analysis parameters to single-cell analysis of two human cell lines. Individual HeLa or K562 cells were processed using the optimized workflow and the resulting peptides were separated using a 5.5 cm μPAC™ HPLC column in a 20-min active gradient. Data acquisition was by DIA with variable window sizes. Data analysis used Spectronaut® with 1% FDR at the protein level and MBR. (**A**): Principal component analysis in Spectronaut®. Each dot represents one cell with HeLa cells shown in red and K562 cells shown in green. (**B**): Bar chart comparison of the average number of identified proteins when analyzing data in the directDIA mode without data enrichment, with data enrichment from 4 replicates each originating from 40 HeLa or 40 K562 cells.

## CONCLUSIONS

We here optimized a sample preparation workflow for single cell proteomics. The most important parameter to minimize losses of such ultra-low input samples seems to perform all steps in a single pot. By omitting any transfer step but inject directly from a 384 well plate, not only proteome coverage but also reproducibility was improved. The wells of a 384 plate thereby serve for all workflow steps from cell isolation to injection into the LC-MS system. Furthermore, preventing drying of samples and the addition of fresh protease after 30 min clearly improved our results. We hypothesize that this not only elongates the total time of efficient digestion but also reduces adsorption of hydrophobic peptides on the well walls. This effect was supported by supplementation of DMSO for sample storage keeping peptides solubilized.

The CellenONE® robot enabled effective and semi-automated cell isolation, served as incubation chamber with controlled temperature and humidity for lysis and digestion, and automatically performed hydration. Although this study was performed using the CellenONE® instrument for all steps except cell isolation, the workflow could be easily implemented without it, since the volumes used are reasonably pipette-able (1 μL) and sample preparation is performed in standard 384-well plates. The 384-well plates are also compatible with alternative cell isolation machines, such as FACS devices, which are available in many labs.

Because a comprehensive workflow includes more than just sample preparation, the complete set of steps, including LC separation, and MS data acquisition, analysis, and interpretation, were also evaluated and optimized. The short μPAC™ HPLC column with its highly ordered and superficially porous brick-shaped pillars provided excellent peptide separation and fast loading and equilibration due to exceedingly low backpressure. While LC-MS analysis in 5-min gradients is possible and generated more than 700 average protein IDs using MS1-only acquisition, best results were obtained using a 20-min active gradient with a total run-to-run time of 39 min. With this cycle time, 37 samples can be analyzed per day. Though this is low throughput compared to multiplexed workflows, multiplexed workflows can suffer from ratio compression and precursor co-isolation which interfere with accurate quantification.^14,25–27^ Because LFQ strategies offer unbiased and accurate quantification, and are fully compatible with DIA, they are preferred when high throughput is not required.

The optimized workflow identified more than 1000 proteins over four orders of magnitude in abundance in a single cell using DDA and data processing without MBR. More than 1790 proteins were identified when data were processed with MBR. Proteomic depth was further improved by using WWA or DIA instead of DDA, providing identification of up to 2250 proteins in a single cell and enabling differentiation of cell types based on protein abundance differences. Easy to implement, robust and sensitive: the optimized workflow has the potential to become the gold standard for studies aiming to dig as deeply as possible into the proteome of individual cells, without the need to use carriers or labelling. In addition, the here developed techniques are not only suitable for single cell applications but in general for all proteomic studies that are limited in input amount requiring loss-free handling and will therefore be highly valuable for a broad scientific community.

## METHODS

### Sample preparation

HeLa cells were cultured at 37°C and 5% CO_2_ in Dulbecco’s Modified Eagle’s Medium (DMEM) supplemented with 10% FBS (10270, Fisher Scientific™, USA), 1x penicillin-streptomycin (P0781-100ML, Sigma Aldrich, Israel) and 100X L-Glut 200 mM (250030-024, Thermo Scientific™, Germany). After trypsinization with 0.05% Trypsin-EDTA (25300-054, Thermo Scientific™, USA) cells are washed 3x with phosphate-buffered-saline (PBS). K562-WT cells were cultured at 37°C and 5% CO_2_ in Gibco Roswell Park Memorial Institute (RPMI -1640) medium (Fisher Scientific™, USA). For harves K562 were pelleted by centrifugation (117g, 4°C) and washed 3x with phosphate-buffered-saline (PBS). All cells were resuspended n PBS at 200 cells/μL for isolation within the CellenONE®.

Benchmarking experiments mimicking single cell digests were prepared from dilutions of the Pierce™ HeLa (#1862824, Thermo Scientific™) protein digest standard in 0.1% TFA with or without 5 % DMSO supplemented as indicated.

Sample lysis and digestion is performed within a clean 384 well plate inside the CellenONE®. 1 μL of a master mix containing 0.2% DDM (D4641-500MG, Sigma Aldrich, Germany), 100mM TEAB (17902-500ML, Fluka Analytical, Switzerland), 3 ng/μL trypsin (Trypsin Gold, V5280, Promega, USA), 0.01 % enhancer (ProteaseMAX™, V2071, Promega, USA) is predispensed into the wells. To limit evaporation humidity is set to the 85%. Individual cells at 18 - 25 μm diameter and a max elongation of 1.5 are sorted by the CellenONE® into the respective wells. This is followed by incubation for 2 h at 50 °C at 85% relative humidity inside the instrument. Samples were kept hydrated every 15 min by automated addition of 500 nL water to each well. After 30 min of incubation, additional 500nL of 3ng/μL trypsin are added which replaces one hydration step. After lysis and digestion, 3.5 μL of 0.1% TFA with 5% DMSO were added to the respective wells for quenching and storage. Samples were either directly injected from the 384 well plate or transferred to a PEG coated PCR tubes (PCR-02-L-C, Axygen®) or to the autosampler glass vials (601801655, Thermo Scientific).

### LC-MS/MS analysis

Samples were analyzed using the Ultimate 3000 RLS-nano high-performance liquid chromatography (Thermo Scientific™) or the Vanquish™ Neo UHPLC system (VN-S10-A01, Thermo Scientific™). Peptides were separated on a 5.5 cm brick shape pillar column prototype (μPACTM, Thermo Fisher) or on a nanoEase M/Z Peptide CSH C18 column (130Å, 1.7 μm, 75 μm x250 mm, 18600810, Waters, Germany) as indicated in a trap and elute setting using an Acclaim™ PepMap™ 100 C18 trapping column (5 μM, 0.3 mm x 5 mm, Thermo Scientific™). All columns were operated at 50°C and connected to an EASY-Spray™ bullet emitter (10 μm ID, ES993; Thermo Fisher Scientific) An electrospray voltage of 2.4 kV was applied at the integrated liquid junction of EASY-Spray™ emitter. Mass spectrometry measurement was performed by an Orbitrap Exploris™ 480 Mass Spectrometer (Thermo Scientific™) equipped with a FAIMS electrode (Thermo Scientific™).

If not otherwise noted, peptide separation was performed at a constant flow rate of 250 nL/min using the following 20-min active gradient: 1% buffer B (acetonitrile (ACN) with 0.08% formic acid) and 99% buffer A (0.1% formic acid (FA)) from minute 0 to 2.5, rising to 2.5% B from minute 2.5 to 2.75, continuing to rise to 25% until minute 17, followed by an increase from 25 to 40% in the last 5 min. The column was washed by increasing buffer B to 97.5% until minute 22 for 10 min before buffer B concentration was decreased back to 1% for re-equilibration. The gradient was shortened to 10 and 5 min for the bench-marking studies. When using the packed-bed column, re-equilibration was performed for 18 min at 250 nL/min. The μPAC™ HPLC column was used on the Vanquish Neo HPLC System with fast loading and equilibration at a maximum pressure of 350 bar and a maximum flowrate of 1 μL/min. For MS1-only acquisition, a linear gradient was applied that ranged from 1 to 40% buffer B only but with no other changes compared to the above-described gradient.

The mass spectrometer and FAIMS Pro Interface parameters are provided in Supplemental Table 1 and Supplemental Table 2.

### Data analysis

If not otherwise noted, data analysis was performed using CHIMERYS™ as a Proteome Discoverer software node (v3.00.757, Thermo Scientific). Data were searched against the human reference database from Uniprot (version: 20.08.2021 UP000005640, 20300 sequences, 11359367 residues) and a customized contaminants database (364 sequences; 142459 residues). For evaluation of the FDR a yeast database was used (version 22.12.2022 UP000002311, 6050 sequences). Trypsin was specified as proteolytic enzyme, cleaving after lysine (K) and arginine (R) except when followed by proline (P) and up to two missed cleavages were allowed. Fragment mass tolerance was limited to 20 ppm and carbamidomethylation of cysteine (C) was set as a fixed modification and oxidation of methionine (M) as a variable modification. Identified spectra were rescored using Percolator^28^ and results were filtered for 1% FDR on peptide and protein level. Abundance of identified peptides was determined by LFQ using IMP-apQuant^22^.

For benchmarking purposes, data were also analyzed using MSAmanda (v2.0.0.19742)^21^, SpectroMine™ (Version 3.2.220222.52329, Biognosys AG), MSFragger^23^, and IonQuant^24^ in FragPipe (Version 18.) For DIA data processing, Spectronaut™ (Version 16.1.220730.53000, Biognosys AG) was used in the directDIA mode. For MS1 only, raw files were converted to mzML files using MSConvert^29^ (v 3.0.21084) enabling the option Peak Picking. The resulting files were analyzed using ms1searchpy^19,20^. Detailed search settings can be found in Supplemental Table 3.

PCA plots were generated using the respective post-analysis function within Spectronaut. The dendrogram was generated using hierarchical clustering (R base function hclust) and the library dendextend. Pairwise sample distance is based on the correlation of log transformed abundances subtracted from one (d = 1-corr(log(a))) with d = distance, a = abundance matrix.

## ASSOCIATED CONTENT

### Data availability

All raw and result files are available for download in the PRIDE repository^30^ using the identifier PXD037719.

### Supporting Information

Supporting information is available to download free of charge.

**Supplemental Figure 1**: Single cell isolation by visual detection using the cellenONE®.

**Supplemental Figure 2**: Venn diagram showing proteins commonly found in a representative replicate from 250 pg HeLa

**Supplemental Figure 3**: Reachable dynamic range from one cell, label free and using DDA

**Supplemental Figure 4**: Improving recovery and reproducibility by tuning workflow parameters

**Supplemental Figure 5**: Dendrogram showing differentiation of cell types from 40 x cell input and no cell controls

**Supplemental Table 1**: Label free MS/MS method for DDA, DIA and MS1only

**Supplemental Table 2**: Summary of the variable isolation windows used for DIA

**Supplemental Table 3:** Summary of data analysis settings for different software and search algorithms

## AUTHOR INFORMATION

### Author Contributions

EM and MM performed experiments, data analysis and wrote the manuscript. EM and MM contributed equally to this work. MM and KM conceptualized the study. GD helped with data analysis and R scripting.

## ACKNOWLEDGMENT

This work funded by the EPIC-XS, Project Number 823839, by the Horizon 2020 Program of the European Union, by the project LS20-079 of the Vienna Science and Technology Fund and the by the ERA-CAPS I 3686, P35045-B, P32054 (FB) and P33380 (FB) project of the Austrian Science Fund. We thank the IMP for general funding and access to infrastructure and especially Karel Stejskal and Gabriela Krššáková, for their continuous laboratory support, Fränze Müller for help in setting up DIA methods and Cynthia Smith, Julian Saba and Rupert Mayer for proofreading the manuscript. Our deepest appreciation also goes to Mark Ivanov and Julia Bubis for their effective and fast support in setting up ms1searchpy and their help in troubleshooting. Special gratitude goes to Ulla Schellhaas for providing us with K562 cells.

