## Supplemental Figures and Tables for "Robust and easy-to-use one pot workflow for label free single cell proteomics"

### Supplementary Figures

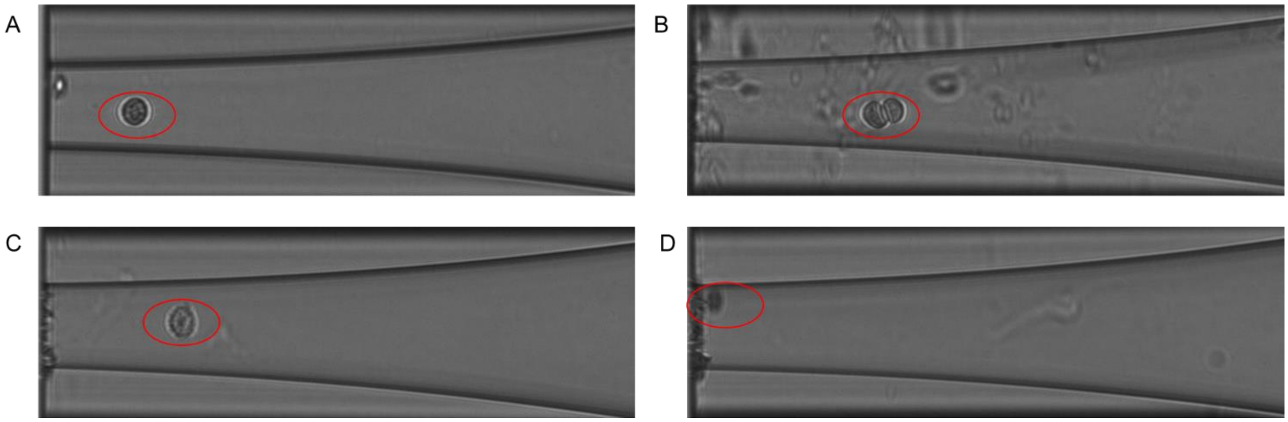


*Supplemental Figure 1:* ***Single cell isolation by visual detection using the cellenONE®*** *(**A**): One intact single HeLa cell is isolated. (**B**): Two cells are co-isolated by accident. (**C**): A single cell whose membrane looks very uneven and that might be inviable. (**D**): No cell is isolated by accident as a condensation droplet on the outer wall mimics a cell.*

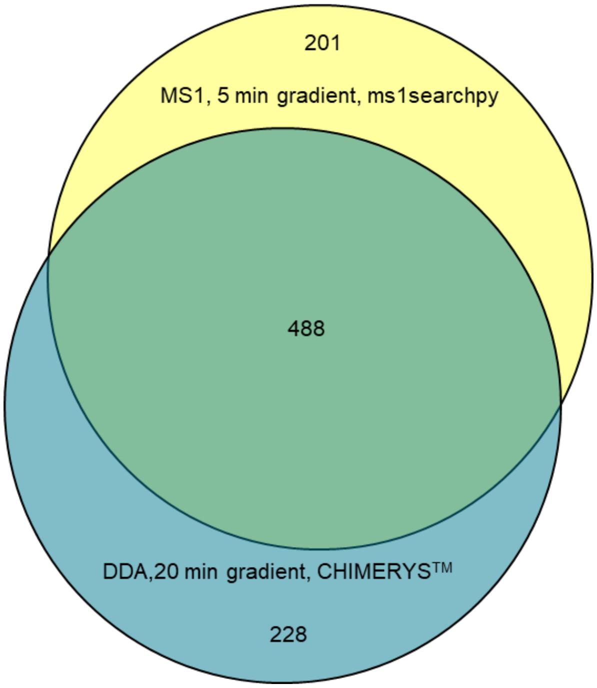


*Supplemental Figure 2:* ***Venn diagram showing proteins commonly found in a representative replicate from 250 pg HeLa.*** *A comparison of the proteins commonly found in a 5-min active gradient with MS1-level spectra used for identification (ms1serchpy) and in a 20-min active gradient, using DDA, and CHIMERYS™ for data analysis.*

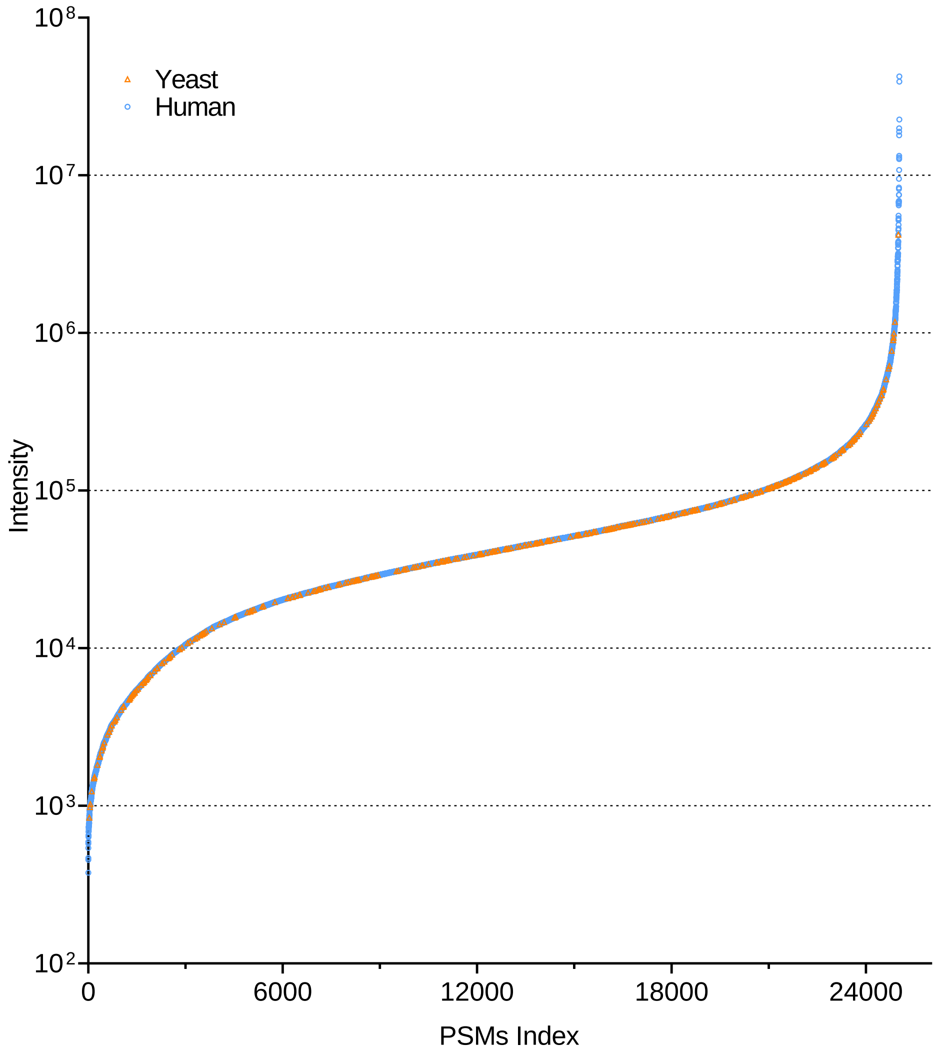


*Supplemental Figure 3:* ***Reachable dynamic range from one cell, label free and using DDA****. Data obtained from a single HeLa cell using the optimized workflow, acquired using a DDA method, and analyzed using CHIMERYS™. PSMs identified from the single-cell run are indexed based on their precursor intensity. PSM matches originating from the yeast proteome database are highlighted in orange.*

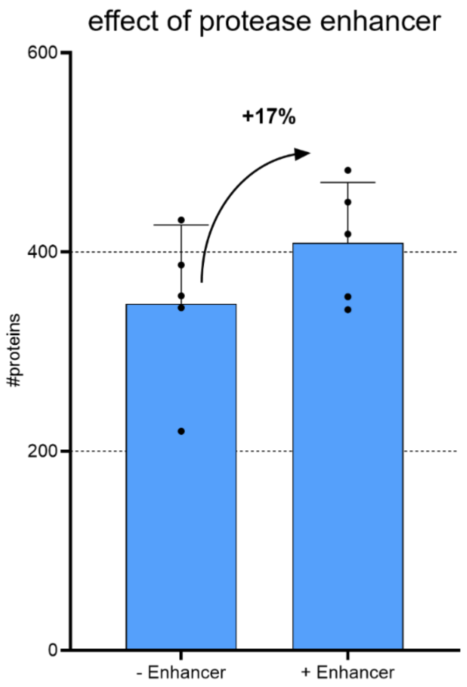


*Supplemental Figure 1:* ***Improving recovery and reproducibility by tuning workflow parameters****. Cells were digested with Trypsin Gold with and without the addition of the protease enhancer ProteaseMAX (Promega)*

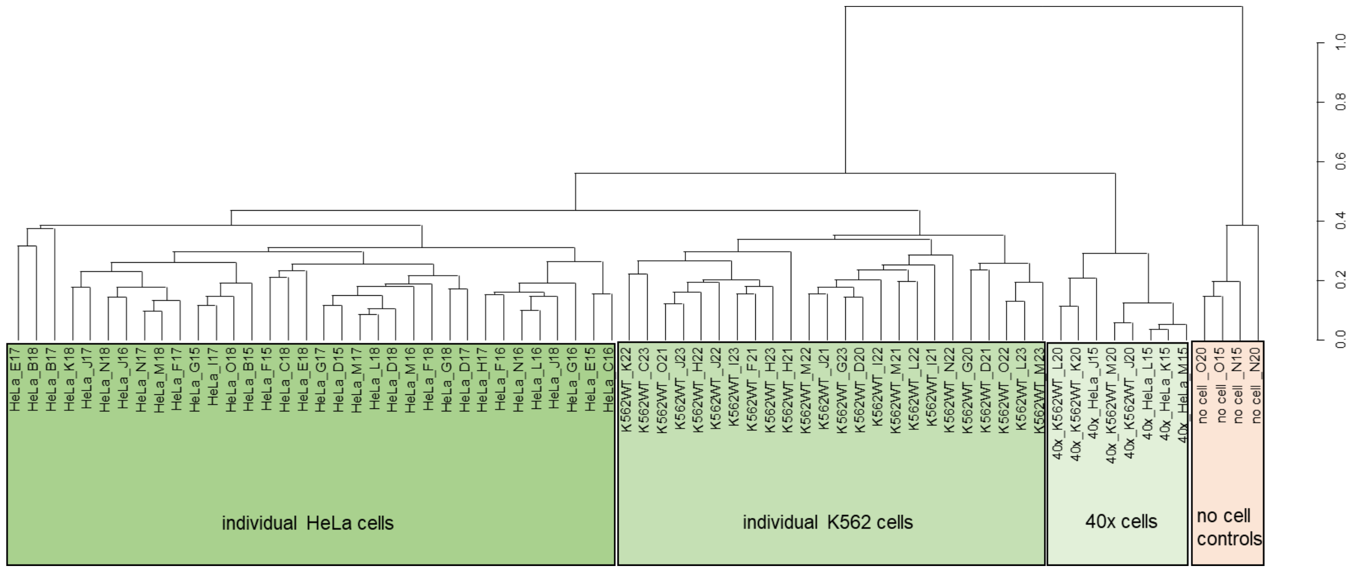


*Supplemental Figure 2:* ***Dendrogram showing differentiation of cell types from 40 x cell input and no cell controls****. Data from Figure 8 but including higher input (40 cells) and no input (no cell) data. No cell controls were processed within the cellenONE in the exact same way as all samples, but no cell was added.*

**Supplementary Tables.**

| Settings | DDA | DIA | MS1only |
|---|---|---|---|
| **Full Scan** | | | |
| *Resolution MS1* | *120000* | 120000 | 120000 |
| *Scan range* | *375-1200* | 375-1200 | 375-1200 |
| *FAIMS CV* | *-50* | -50 | -40, -50, -60 |
| *AGC Target (%)* | *300* | 300 | 300 |
| *Maximum Injection time (s)* | *Auto* | Auto | Auto |
| **MS2 scan** | | | |
| *Number of dependent scans* | *10* | | |
| *Minimum precursor intensity* | $5.0*e^{3}$ | | |
| *Charge state* | *2-5* | | |
| *Dynamic exclusion time (s)* | *120* | | |
| *Isolation window (m/z)* | *2* | | |
| *HCD Collision Energy (%)* | *30* | 30 | |
| *Resolution MS2* | *60000* | 60000 | |
| *First mass (m/z)* | *120* | | |
| *AGC Target (%)* | *75* | 75 | |
| *Maximum Injection Time (s)* | *118* | 118 | |

*Supplemental Table 1: Label free MS/MS method for DDA, DIA and MS1only*

| Precursor mass range (m/z) | Isolation window (m/z) | Number of scan events |
|---|---|---|
| 375-500 | 25 | 4 |
| 500-600 | 13 | 7 |
| 600-850 | 8 | 31 |
| 850-900 | 13 | 3 |
| 900-1200 | 25 | 9 |

*Supplemental Table 2: Summary of the variable isolation windows used for DIA*

| | MSAmanda | SpectroMine® | Spectronaut™ | FragPipe | Ms1searchpy |
|---|---|---|---|---|---|
| Proteolytic enzyme | Trypsin, cleavage at K and R if not followed by P | | | | |
| Missed cleavages | max. 2 | | | | |
| Precursor mass tolerance | ±5 ppm | | | | ± 8 ppm |
| Fragment tolerance | 10 ppm | | | | - |
| Minimum peptide length | 6 | | | | 7 |
| Variable modifications | Oxidation at methionine, acetylation at protein N-terminus | | | | Not defined |
| FDR (peptide, protein level) | 1% | | | | |

*Supplemental Table 1 Summary of data analysis settings for different software and search algorithms*
